# Survival strategies of the entomopathogenic nematode encapsulated by nanoparticle-stabilized emulsions at rapid desiccation

**DOI:** 10.1101/2025.07.14.664280

**Authors:** Jayashree Ramakrishnan, Satheeja santhi Velayudhan, Adi Faigenboim, Ahmed Nasser, Mohamed Samara, Eduard Belausov, Victoria Reingold, Amit Horn, Karthik Ananth Mani, Guy Mechrez, Itamar Glazer, David Shapiro-Ilan, Dana Ment

## Abstract

Water retention is essential for survival under desiccating conditions, especially for entomopathogenic nematodes (EPNs) used as biocontrol agents. Foliar application of EPNs causes rapid desiccation (RD), severely reducing their survival and efficacy. We investigated the physiological and molecular responses of *Steinernema carpocapsae* to RD, delivered in a novel nanoparticle-stabilized emulsion to improve tolerance under variable humidity. We hypothesized that the formulation would enhance EPNs survival, by promoting early stress-responsive pathways and water-retention mechanisms. Formulated nematodes exhibited delayed water loss and increased survival under low humidity compared with non-formulated controls. The emulsion mitigated RD by enhanced hydration retention and reduced water loss, effects that were strongly associated with differential trehalose accumulation. In addition, maintenance of basement membrane integrity and cytoskeletal organization, emerged as a critical component of desiccation. These findings advance understanding of RD tolerance in EPNs and demonstrate how tailored formulation strategies can protect biocontrol agents under desiccation stress.

## INTRODUCTION

Water is essential for life’s fundamental biological processes. During desiccation, rapid water loss results in significant reductions in cell volume, accompanied by structural modifications and alterations in both extracellular and intracellular solute concentrations^1^. The genetic, biochemical, and molecular machineries that enable survival in extreme environments present a complex interplay of adaptation strategies that influence tolerance to water stress^2^. Insights across groups (bacteria, yeast, plants, tardigrades, nematodes, and insects) suggest both conserved and diverse adaptation strategies to withstand water loss and desiccation^3–5^.

Entomopathogenic nematodes (EPNs) from the families Steinernematidae and Heterorhabditidae are slow-dehydration strategists used commercially for insect biocontrol ^6–9^. EPN sensitivity to abiotic stresses^10,11^, including low tolerance to rapid desiccation (RD) defined as rapid water loss over minutes to hours, severely impacts their viability and hence, biological efficacy^12,13^. Thus, the infective juvenile (IJ) stage of EPNs offers a valuable model to investigate desiccation tolerance due to their ability to survive prolonged dehydration through quiescence, characterized by significantly reduced metabolic activity until favorable conditions return^14^. Several protection strategies have been reported during preconditioning (72 h at high humidity)^7,15,16^. However, studying the innate adaptation strategies employed by EPNs and enhancing their resistance to rapid water loss are critical to improving their persistence and biocontrol efficacy^17–19^.

We recently characterized the inverse relationship between water-loss rates in EPNs at different relative humidities (RHs) and RD^20^. That study revealed distinct differences in membrane-packing abilities during the initial drying period, indicating strong water-binding properties of *Steinernema carpocapsae*. Those observations led to questions about the underlying mechanisms that enable EPNs to withstand rapid water loss. We further found that formulations such as nanoparticle-stabilized Pickering emulsions (titania Pickering emulsion [TPE] and silica Pickering emulsion gel [SPEG]), tailored to reduce water loss, resulted in enhanced survival and efficacy of *S. carpocapsae* IJs^21^. Building on those findings, we hypothesized that EPN survival and efficacy under RD involve distinct changes in physiological, biochemical, and molecular mechanisms. We therefore recruited various strategies to approach the broader question of what changes enable survival under RD. This information could provide deeper insights into the nuances of RD toward integrating intervention strategies that provide maximal protection while maintaining organism functionality during desiccation stress.

## RESULTS

### Effects of physiological water loss-delaying strategies on EPN IJ survival

*In-vitro* survival of formulated and non-formulated *S. carpocapsae* IJs was assessed at 64% RH and 54% RH over 144 h (Fig. 1A). Irrespective of humidity, survival of formulated IJs was significantly higher than that of the non-formulated controls (IJs in water) (Fig. 1B and S1). TPE- and SPEG-treated IJs maintained viability for up to 120 and 144 h, respectively. In contrast, controls reached 0% survival at both 54% RH (F_(14,349)_ = 48.0069, *P* < 0.0001) and 64% RH (F_(14,317)_ = 39.6917, *P* < 0.0001). At 64% RH, survival in the control decreased by 80% within 16 h, whereas formulated IJs maintained >80% viability for up to 72 h. Survival rates for TPE and SPEG IJs were not significantly different for the first 24 h (F_(3,317)_ = 1.000, *P* > 0.05), but diverged significantly after 16 h at 54% RH (F_(4,349)_ = 23.4109, *P <* 0.0001), indicating varying responses in different formulations with humidity level and time. Although SPEG consistently provided higher protection at both humidity levels, increased variability at >72 h suggested destabilization of the emulsion system.

**Fig. 1.**
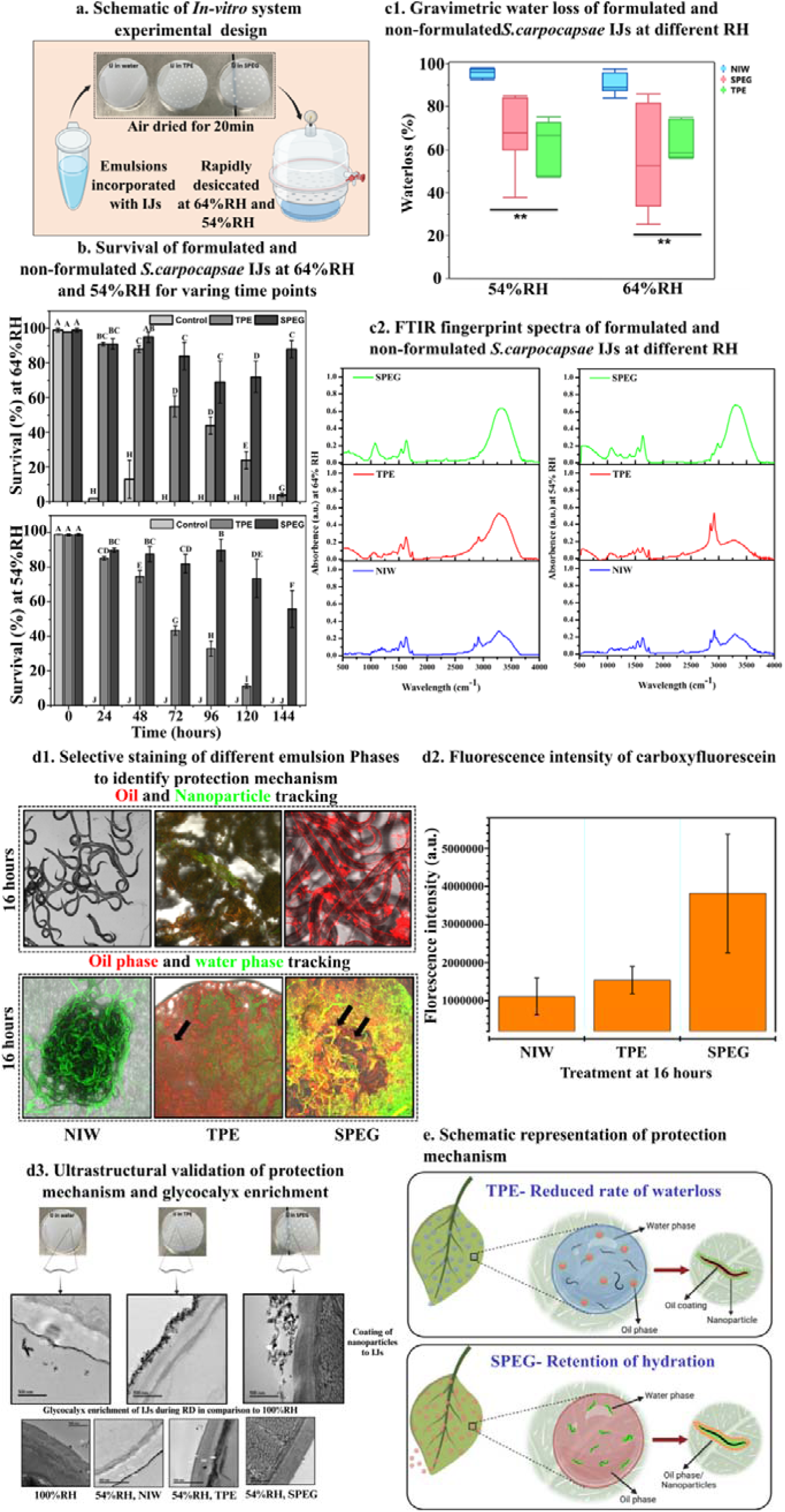
Effect of low RH and formulations on survival and water-loss characteristics of *S. carpocapsae* infective juveniles (IJs) following rapid desiccation (RD). (**A**) Schematic illustration of the experimental design: non-formulated (in water) and formulated (titania Pickering emulsion [TPE] and silica Pickering emulsion gel [SPEG]) IJs were incubated in desiccators at 64 ± 5% or 54 ± 5% RH (23°C) for RD, then transferred to 55-mm petri plates with 7–10 ml distilled water and incubated overnight for viability determination. (**B**) IJ (formulated and non-formulated control) survival was determined at 0–144 h post-RD at both RHs. Error bars represent standard error. (**C1**) Gravimetric water loss in IJs after RD exposure for 24 h at 64% RH or 54% RH. (**C2**) FTIR spectral fingerprint region of *S. carpocapsae* IJs post-RD for 72 h at 64% or 54% RH and after subtraction from respective negative emulsion-only controls. (**D1**) Confocal microscopy images of formulated and non-formulated IJs: nematodes in water (NIW), TPE or SPEG were drop-casted on slides and rapidly desiccated at 54% RH for 16 h. Emulsion oil phase is labeled with Nile red, and nanoparticles are labeled with carboxyfluorescein (upper panel). Modified procedure with carboxyfluorescein-labeling of water phase (lower panel). Visualization of oil-phase and nanoparticle tracking (upper panel), and oil-phase and water-phase tracking (lower panel), highlights localization and protective interactions of the formulations. (**D2**) Fluorescence intensity of carboxyfluorescein, used as an indicator for water in Pickering emulsions after RD for 16 h at 54% RH. (**D3**) Ultrastructural validation of the protective mechanisms (upper panel) and glycocalyx enrichment in rapidly desiccated SPEG and NIW compared to controls maintained at 100% RH (lower panel). (**E**) Schematic representation summarizing the protective mechanisms conferred by TPE (upper panel) and SPEG (lower panel) against water loss under RD conditions.

### Effects of RH and formulations on water loss in IJs

Gravimetric analysis was conducted on formulated vs. non-formulated IJs for 24 h (Fig. 1C1). At 64% RH and 54% RH, the control IJs lost approximately 98% of their water content, compared to 50 and 70% water loss in SPEG and TPE IJs, respectively. Although no significant differences were observed in the rate of water loss between 54% RH and 64% RH (F_(1,26)_ = 1.3032, *P* > 0.05), formulated IJs exhibited significantly less water loss compared to controls (F_(2,26.1)_ = 19.3396, *P* < 0.001), demonstrating the formulations’ effective protection by significantly slowing IJs’ rate of water loss. Measurements were not conducted after 24 h due to negative values obtained upon subtraction from controls (emulsion only) (Fig. S2).

The Fourier transform infrared (FTIR) spectral fingerprint regions for EPN IJs in water, TPE, and SPEG varied in response to desiccation at 64% RH and 54% RH for 72 h (Fig. 1C2). Although the entire FTIR spectrum is required to reliably predict IJ water content ^20^, Pickering emulsions complicate spectral interpretation due to overlapping regions of IR-active components/phases. We therefore focused specifically on the 3000–3500 cm^-1^ region, corresponding primarily to hydroxyl (OH)-bond vibrations, to reliably assess hydration retention in formulated EPNs. Control IJ samples showed the lowest peak intensity compared to TPE- and SPEG-formulated IJs after 72 h of desiccation. SPEG-formulated IJs exhibited the highest peak intensity, indicating better OH-bond-retention capacity under reduced humidity. At 54% RH, a noticeable decrease in OH-bond peak intensity was observed in TPE-formulated IJs, corresponding to their significantly lower survival compared to SPEG-formulated IJs (F_(1,349.1)_ = 65.8327, *P* < 0.0001).

### Physiological protection mechanism imparted by emulsions for EPN survival

To elucidate the protective mechanism of Pickering emulsions and identify the roles of their components, we labeled the oil phase and titanium dioxide nanoparticles using Nile red and 5,6-carboxyfluorescein, respectively, through surface modification via amidation (EDC chemistry)^22^. The SPEG silica nanoparticles (commercial hydrophobic particles AEROSIL® R 972) were not labeled due to the absence of functional groups required for surface modification.

Microscopy images revealed adhesion of the titanium nanoparticles to the TPE-formulated IJ cuticles, presumably through physical adsorption or attachment (Fig.1D1, upper panel). In contrast, control IJs suspended in water exhibited a shrunken and dried appearance at 16 h. A distinct thin red coating on formulated EPN cuticles indicated the presence of oil (Fig. 1D1, upper panel, right). Similar behavior was previously noted with SPEG, where the hydrophobic nature of nanoparticles within the inverse water-in-oil (W/O) emulsion resulted in analogous red-labeling patterns^23^. Notably, this protective coating appeared only after RD, indicating its IJ-protection specificity (Fig. S3). To determine whether formulations retain hydration after desiccation, the aqueous and oil phases were labeled with carboxyfluorescein and Nile Red, respectively. A layer of carboxyfluorescein was clearly deposited on the control IJs (Fig. 1D1, bottom panel). As previously observed with TPE, IJs were coated with oil and titanium nanoparticles. In contrast, a distinct water/polymer layer was observed surrounding the SPEG-formulated IJs (Fig. 1D1, bottom panel, right). Fluorescence intensity measurements (Fig. 1D2) confirmed higher carboxyfluorescein retention for SPEG (3 × 10^6^ fluorescence intensity) compared to control (1 × 10^6^) and TPE (1.2 × 10^6^) IJs, further strengthening our findings.

Moreover, SPEG-formulated IJs retained motility after 16 h of RD at 54% RH (Supplementary Video). Collectively, these findings indicated the distinct protective mechanisms of each formulation: SPEG promotes survival through hydration retention, whereas TPE primarily reduces the rate of water loss in IJs during RD (Fig. 1E). Our transmission electron microscopy (TEM) results consistently demonstrated glycocalyx enrichment in SPEG-treated samples (Fig. 1D3). Interestingly, we observed a very dark epicuticle layer in rapidly desiccated IJs compared to controls (100% RH), which could be due to increased epicuticle thickness or glycoprotein accumulation. In addition, glycocalyx enrichment was apparent in rapidly desiccated TPE-treated IJs, though visualization was inconsistent and unclear due to the coating of titanium nanoparticles.

### Core and novel pathways involved in RD tolerance revealed by functional enrichment

Functional enrichment analysis revealed an overrepresentation of distinct biological processes (by gene ontology [GO]) and pathways (by Kyoto Encyclopedia of Genes and Genomes [KEGG]) within the combined clusters (Fig. 2C). Cluster 1,5 encompassed 488 genes that were downregulated compared to controls yet maintained relatively higher expression levels under lower vs. higher humidity (Fig. 2C1). Those enriched at lower humidity included HSP binding (GO:0031072), endopeptidase activity, G-protein-coupled receptor activity (GO:0004930), membrane components (GO:0016020), structural constituent of cuticle (GO:0042302), extracellular region (GO:0005576), and processes related to signal transduction (GO:0007165).

**Fig. 2.**
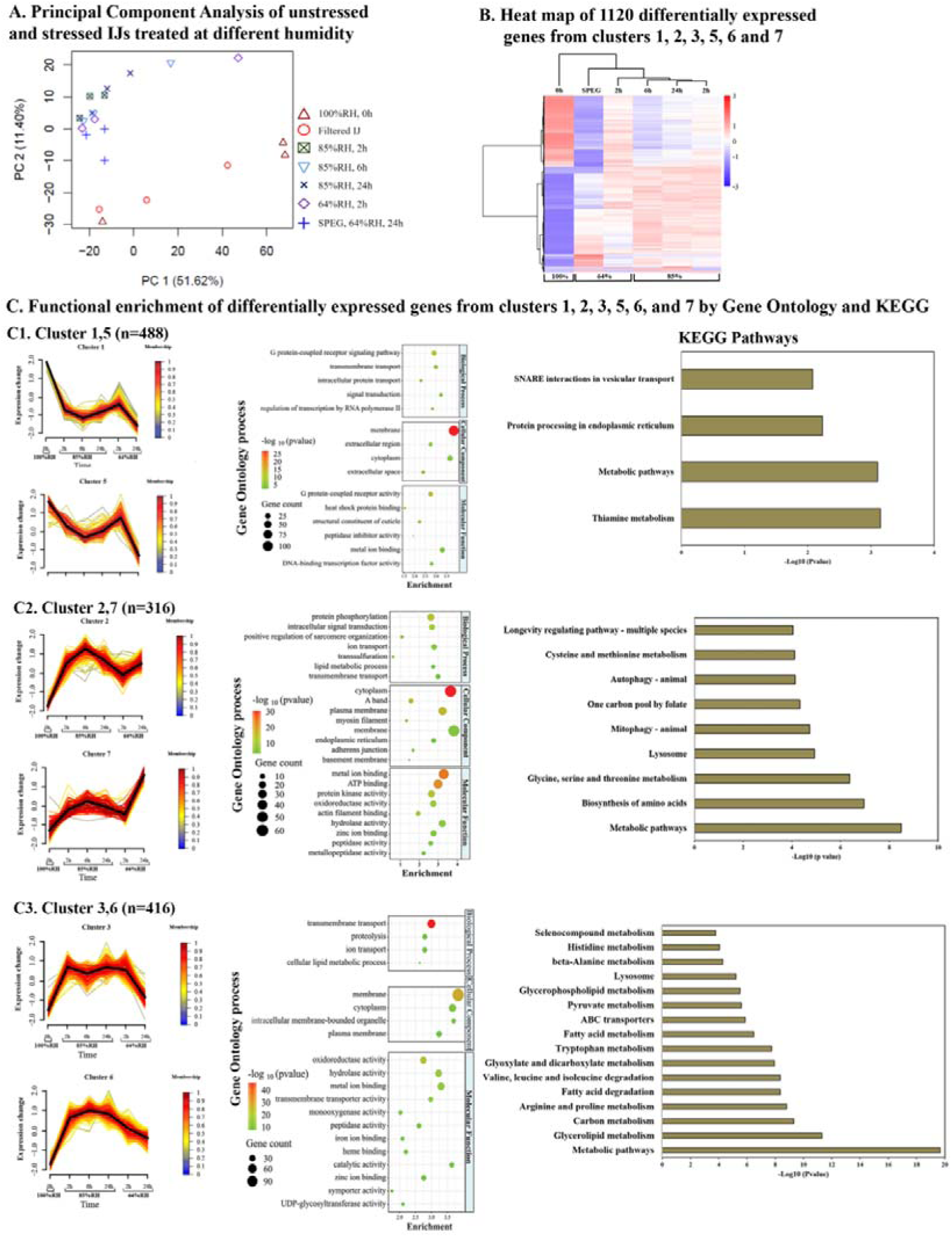
Quality assessment and preliminary screening of transcriptome data. a PCA depicting the clustering patterns of stressed, formulated and control IJs. b Heatmap illustrating the differentially expressed genes (DEGs) in stressed vs. unstressed *S. carpocapsae* IJs. c Hierarchical clustering and functional enrichment analysis using KOBAS databases groups DEGs into clusters based on correlations in expression trends over the experimental time course. c1, Cluster 1,5; c2, Cluster 2,7; c3, Cluster 3,6

Cluster 2,7 consisted of 316 genes that were significantly upregulated compared to the control group (Fig. 2C2). Their expression trends varied across different time points and treatments, with the highest expression observed in formulated treatments (SPEG) and at higher humidity levels compared to lower ones. This cluster likely reflects adaptive changes promoting survival during RD over time. Notably, enriched GO categories related to cellular components and biological processes, such as structural constituent of muscle (GO:0008307), A band (GO:0031672) and M band (GO:0031430), cytoplasm (GO:0005737), and sarcomere organization (GO:0060298). KEGG pathway analysis identified significant upregulation of autophagy-related processes. Notably, our results indicate that membrane organization and sarcomere assembly may represent a critical remodeling process, adjunct to cytoskeletal remodeling^34^, which underlies desiccation tolerance.

Cluster 3,6 comprised 416 genes that were significantly upregulated compared to the control group (Fig. 2C3). A consistent expression trend across humidity levels indicated the fundamental role of these genes in desiccation tolerance. Enriched biological processes within these clusters included transmembrane transport (GO:0055085), cell volume homeostasis (GO:0006884), UDP-glycosyltransferase activity (GO:0008194), hydrolase (GO:0016787) and oxidoreductase (GO:0016491). In addition, membrane-, cytoplasm- and proteolysis-related processes (GO:0016020, GO:0005737 and GO:0006508, respectively) were significantly enriched. KEGG pathway analysis confirmed the involvement of established pathways associated with desiccation or anhydrobiosis, including carbon metabolism, fatty acid metabolism and carboxylate/dicarboxylate metabolism, reinforcing their critical role in desiccation adaptation.

### Transcriptomic evidence suggests that SPEG formulation protects IJs via environmental buffering

A transcriptomic comparison of upregulated genes between desiccated IJs at high humidity (85% RH, all time points) vs. formulated IJs (at low humidity) indicated 2024 (51.23%) genes shared among all samples (Fig. S6, Table S3). The next largest shared cluster included genes that were common to SPEG and the 6 h treatment at 85% RH (721 genes, 18.00%). DEGs unique to SPEG (361 genes, 9.14%) were significantly upregulated with enriched pathways including mucin-O-type glycan biosynthesis (Cel00512, enrichment ratio [ER]: 0.27, log *P* = 2.68) and ECM–receptor interaction (Cel004512, ER: 0.375, log *P* = 2.93). Significantly enriched cellular component GO terms included transferase activity of glycosyl groups (GO:0016757, log *P* = 4.84), basement membrane (BM) (GO:0005604, log *P* = 6.02), extracellular matrix organization (GO:0030198, log *P* = 4.34), ecdysis, collagen and cuticulin-based cuticle (GO:0042395, log *P* = 3.31), peptidyl-serine phosphorylation (GO:0018105, log *P* = 12.47), chromosome (GO:0005694, log *P* = 10.83) and protein folding (GO:0006457, log *P* = 5.27).

Conversely, downregulated processes uniquely associated with SPEG compared to high-humidity treatments were G protein-coupled receptor signaling pathway (GO:0007186; log *P* = 3.00), cytoplasm (GO:0005737; log *P* = 6.22), membrane (GO:0016020, log *P* = 18.60), and integral component of membrane (GO:0016021, log *P* = 16.19).

### RD mirrors osmotic rather than evaporative desiccation processes

To determine fundamental differences between rapidly desiccated (*S. carpocapsae* IJs) and slow-desiccated/preconditioned IJs, we utilized our previous data from preconditioned *Heterorhabditis bacteriophora* subjected to osmotic (20% polyethylene glycol [PEG], 36 h) or evaporative (97% RH, 24 h) desiccation.

Functional enrichment analysis identified significantly enriched GO processes and KEGG pathways (both upregulated and downregulated); these were compared via Venn diagram differentiation among osmotic, evaporative and RD treatments (Fig. 3). Among the upregulated processes, 58% (749 processes) were unique to RD, and 25% (337 processes) overlapped between osmotic desiccation and RD. Notably, only 1.4% of the processes were present in all stress conditions. These findings suggest that IJs encounter significant osmotic stress early on in RD, potentially due to rapidly changing solute concentrations driven by the paracellular movement of water. Unique and shared terms for the RD, osmotic desiccation (PEG) and evaporative desiccation groups are presented in rectangular boxes in Fig. 3. Enriched terms encompassed BM (GO:0005604), ECM (GO:0031012), adherens junction (GO:0005912), hemidesmosome (GO:0030056), sarcomere (GO:0030017) and cytoskeleton (GO:0005856) organization, regulation of cell size (GO: 0008361) and integrin-mediated signaling pathways (GO:0007229). The BM enrichment observed during osmotic desiccation was not significant.

**Fig. 3.**
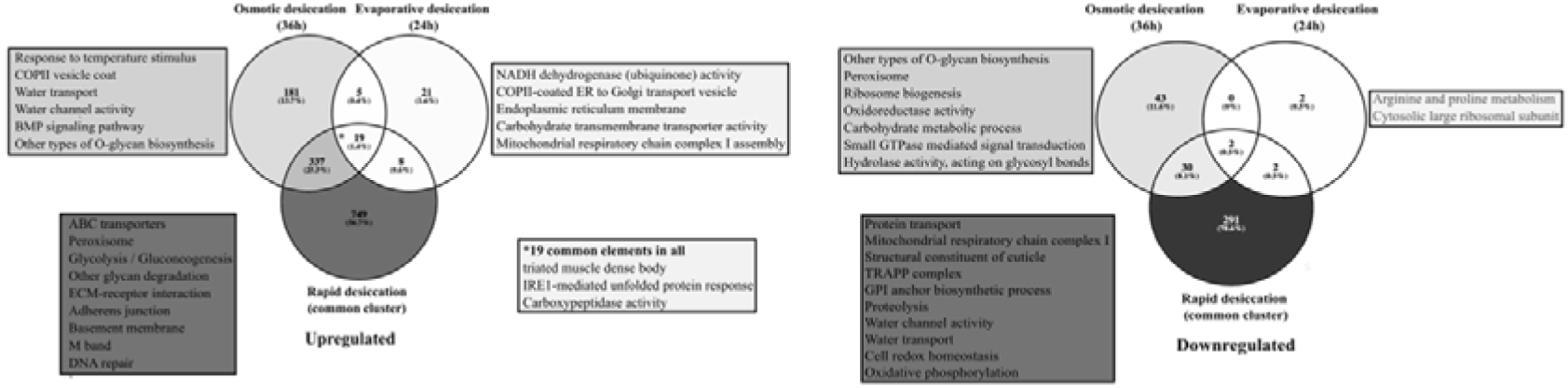
Venn diagram illustrating unique and shared enriched GO processes and KEGG pathways for the differentially expressed transcripts (upregulated and downregulated) of IJs that underwent RD, and preconditioned IJs subjected to osmotic desiccation or evaporative desiccation.

Comparisons of pathways indicated that the Purine metabolism and protein-export pathways were shared between the evaporative desiccation and RD groups. Unique pathways identified in the RD group, compared to previously published Steinernema data ^16^, included ECM–receptor interaction and DNA damage-response pathways, such as the Fanconi anemia pathway, nucleotide excision repair, and mismatch repair. Analysis of downregulated processes revealed suppression of water-channel activity, proteolysis and oxidative phosphorylation (cytochrome I to IV) during RD. Interestingly, osmotic desiccation showed upregulation of water transport, water-channel activity and cytochrome I, whereas cytochrome V (ATP synthase) and water transport were downregulated specifically during RD. These observations suggest that RD imposes significant metabolic and energetic demands on EPN IJs. Commonly enriched terms among all desiccation treatments included the unfolded protein response (UPR) mediated by the endoplasmic reticulum and structural components, such as striated muscle dense bodies (Z-disc), emphasizing the critical roles of protein refolding and stress-signaling hubs in initiating downstream activation of stress effector proteins.

### Gluconeogenesis and the trehalose-biosynthesis pathway

Many anhydrobiotic organisms biosynthesize and accumulate trehalose during preconditioning^24–26^. We thus analyzed gene expression in the trehalose-biosynthesis pathway. The glyoxylate shunt pathway was significantly upregulated in RD IJs vs. controls (ER: 0.42, log *P* = -8.457, Fig. 4A), consistent with trehalose accumulation^27,28^. Gene-expression analyses showed significant differential expression patterns for trehalose-biosynthesis genes (*tps-1, tps-2, icl-1*) under varying humidities and in formulated treatments compared to controls at 100% RH (Fig. 4B). Notably, ICL-1, responsible for glyoxylate shunt, was highly upregulated (logFC: 6.9) under low humidity compared to SPEG treatment (logFC: 5.3). TPS-1 and TPS-2, known for trehalose synthesis, exhibited higher expression (logFC: 3.0) at low humidity compared to formulated SPEG IJs (logFC: 1.8), suggesting trehalose’s central role in protecting membranes and managing bioenergetics during dehydration and rehydration in RD.

**Fig. 4.**
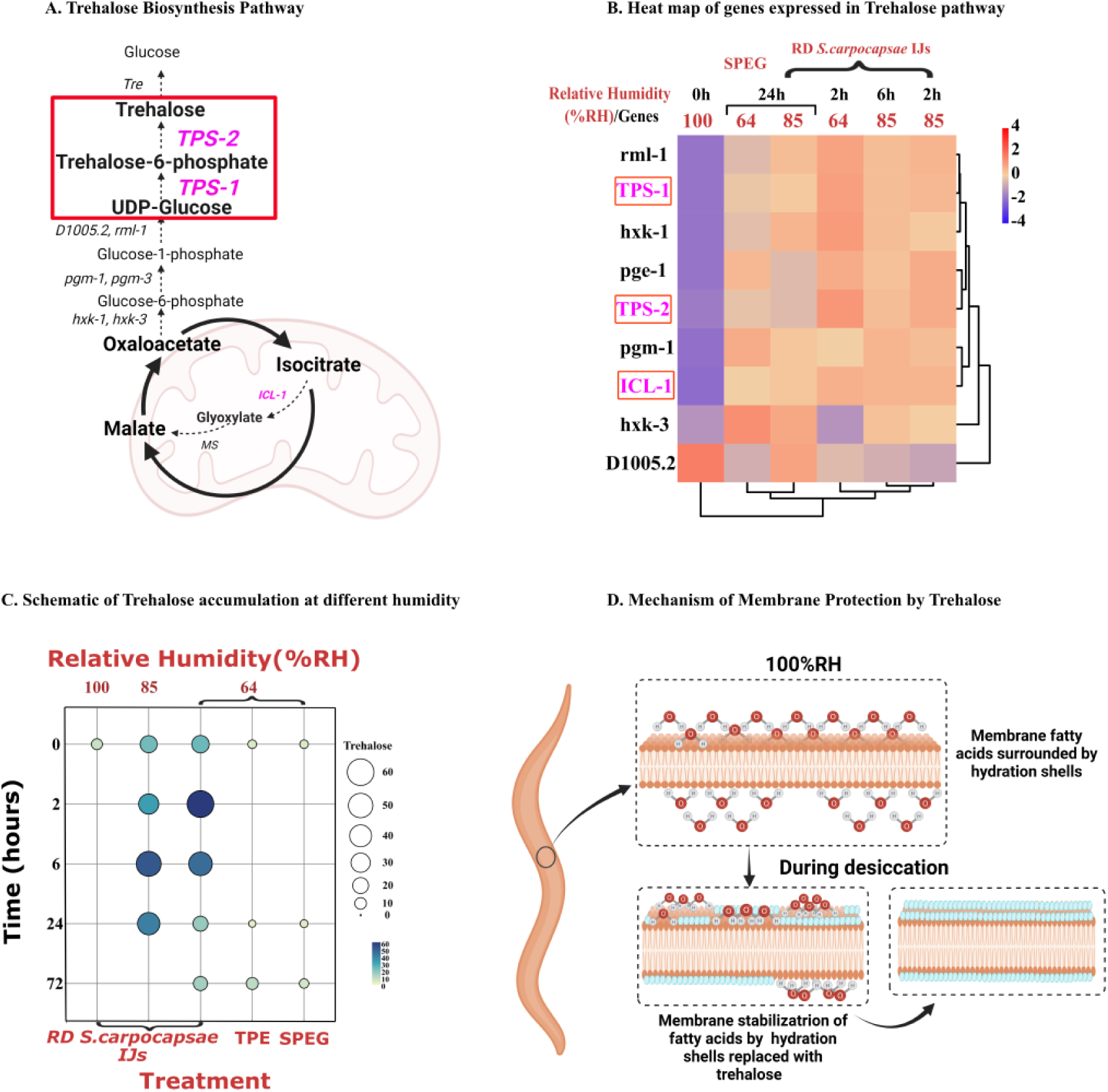
Mechanism of trehalose accumulation in *S. carpocapsae* IJs under RD. (**A**) Trehalose biosynthesis pathway initiated through glyoxylate shunt metabolism, highlighting key genes involved. (**B**) Heatmap of genes expressed in the pathway and their expression patterns under different RH conditions and at various time points. (**C**) Schematic representation of trehalose accumulation (ug/mg) across treatments, environmental conditions and time points. Bubble size and color indicate intensity of accumulation. Experimental design for validating trehalose accumulation: *S. carpocapsae* IJs at 85% and 64% RH and formulated IJs (64% and 54% RH) were rapidly desiccated for 0, 2, 6, 24 and 72 h. Comparison of trehalose accumulation between formulated and non-formulated IJs at 64% RH and 54%RH and *S. carpocapsae* IJs desiccated at different humidity levels (85% RH and 64% RH), measured on a dry weight basis after 72 h of RD. **(D)** Proposed mechanism of trehalose accumulation in IJs during desiccation based on water-replacement hypothesis.

To validate the gene-expression findings, trehalose levels for different samples were quantified by UHPLC with a carbohydrate-specific column. Trehalose accumulation varied significantly among treatments and conditions (Fig. 4C). Initially, filtered *S. carpocapsae* IJs had significantly higher trehalose levels than non-formulated controls (F_(2,14)_ = 7.28732, *P* = 0.0068), aligning with previous studies indicating constitutive trehalose accumulation in *Steinernema* species under stress^16,29^. Although trehalose levels increased over time at higher humidity, this was not statistically significant (F_(3,19)_ = 2.2363, *P* > 0.05). Conversely, under low humidity, trehalose accumulation increased significantly with time (F_(3,17.2)_ = 8.6373, *P* = 0.0010). At 64% RH, trehalose levels were threefold higher within 2 h than in controls (F_(1,17)_ = 24.46, *P <* 0.0001) and 85% RH (F_(1,37)_ = 15.7582, *P* < 0.0003). However, after 2 h, accumulation levels stabilized with no significant differences among humidity conditions. It is also important to note that, compared with preconditioned *S. carpocapsae*, the RD IJs showed significantly higher trehalose accumulation at 85% and 64% RH during early time points (2-6hr) (Data not shown).

Comparisons between formulated and non-formulated IJs showed significant differences in trehalose accumulation across humidity conditions (F_(4,70)_ = 5.532, *P* = 0.0006, Fig. 4C). Control (100% RH) trehalose levels at 0 h were not significantly different from those of formulated samples (F_(2,70)_ = 0.216, *P* > 0.05). Overall, irrespective of humidity (except the controls), trehalose accumulation was highest in desiccated controls, intermediate in TPE IJs, and lowest in SPEG IJs (F_(2,70)_ = 13.362, *P* < 0.0001). Unlike previous findings, trehalose accumulation in desiccated controls was significantly lower at 54% RH compared to 64% RH (F_(1,70)_ = 4.168, *P* = 0.0451).

### Membrane organization and ECM remodeling via BM assembly as critical factors for desiccation tolerance

Enriched GO biological processes identified during RD included cytoskeletal reorganization, actin dynamics, cell volume homeostasis and KEGG pathways such as ECM–receptor interaction. Comprehensive data mining revealed differences in gene expression related to cytoskeleton, sarcomere, and ECM organization, which were visualized using heatmaps (Fig. 5D). Notably, BM-related genes exhibited differential expression across humidity conditions and in the SPEG formulation.

**Fig. 5.**
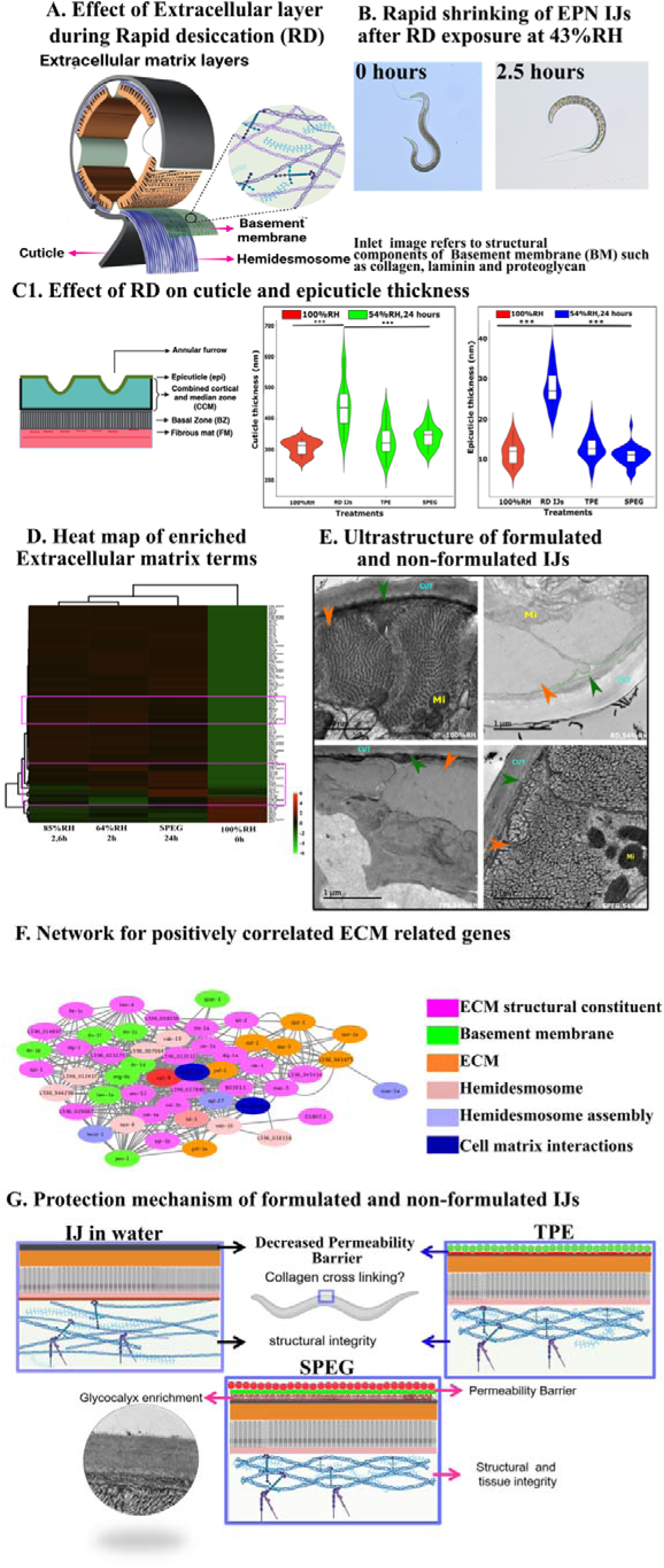
Impact of membrane organization and ECM remodeling of *S. carpocapsae* IJs during RD assessed by ultrastructural microscopy. (**A**) Transverse section of nematodes illustrating ECM layers (cuticle, hypodermis/hemidesmosome and basement membrane [BM]); image courtesy Wormbook. (**B**) Rapid shrinking of *S. carpocapsae* IJs after 2.5 h exposure to 43% RH. (**C**) Effect of RD on cuticle thickness, especially the epicuticle (calculated from cryo-TEM images) (**D**) Heatmap of enriched processes, such as BM, hemidesmosome, focal adhesion and ECM. (**E**) Cryo-TEM images of desiccated IJs after 24 h at 54% RH. Treatments include control (100% RH), rapidly desiccated IJs and formulated IJs (TPE and SPEG). Green arrowheads indicate BM; orange arrowheads indicate sarcomere assembly. (**F**) Network analysis of genes associated with BM, ECM and hemidesmosome (HD) positively correlated to RD. Different colors represent distinct GO terms. (**G**) Proposed protective mechanisms involved in control vs. formulated IJs.

To validate these transcriptomic findings, cryo-TEM imaging was used to compare the membrane organization in rapidly desiccated IJs (formulated and non-formulated) at low humidity (Fig. 5E). Sarcomere organization was severely disrupted in all RD treatments compared to controls (100% RH). Membrane organization was particularly disrupted in rapidly desiccated IJs treated with water and TPE, while notably preserved in SPEG-treated IJs. This suggests that membrane structural plasticity has a significant influence on nematode survival under abiotic stress conditions. In addition, we observed differences in BM structural integrity. Rapidly desiccated IJs exhibited extensive damage below the dense Z-disc layer along the circumferential axis, whereas TPE- and SPEG-treated IJs retained intact ECM structures. This aligns with the survival data, reinforcing the importance of intact ECM for nematode survival, while tissue integrity and membrane organization are essential for IJ efficacy. Further analysis explored changes within the ECM layers, specifically targeting the cuticle. Genes associated with structural components of the cuticle, their maintenance, turnover and synthesis were predominantly downregulated, as reported previously^30^. However, genes involved in cuticle crosslinking were upregulated (Fig. S7). Cuticle thickness across layers (epicuticle, cortical and median layers combined, basal layer, hypodermis) was quantitatively measured via cryo-TEM (Fig. 5C1). Treatments significantly influenced overall cuticle thickness (F_(3,23)_ = 6.6008, *P* = 0.0028), which was significantly greater in rapidly desiccated IJs compared to TPE-formulated IJs (*P =* 0.00081; 95% CI: 284.45, 384.15), SPEG IJs (*P* = 0.0166; 95% CI: 311.26, 375.98) and controls (100% RH, *P* = 0.0116; 95% CI: 246.79, 372.26). No significant differences were detected among TPE (*P* = 0.9099), SPEG (*P* = 0.7987) and control groups, suggesting a protective effect of the emulsions.

Epicuticle thickness varied significantly among treatments (F_(3,23)_ = 49.5867, *P* < 0.0001), with rapidly desiccated IJs showing a 154% increase compared to controls (100% RH, *P* < 0.0001; 95% CI: 3.907, 19.35), TPE (*P* < 0.0001; 95% CI: 10.603, 15.287) and SPEG (*P* < 0.0001; 95% CI: 9.133, 12.216). Furthermore, the TPE and SPEG groups did not differ significantly from the controls (100% RH, P > 0.05), emphasizing that epicuticle thickness modulation is critical for maintaining a physiological barrier against water loss, consistent with previous studies ^46, 47^.

## DISCUSSION

We investigated the physiological, molecular and biochemical strategies underlying *S. carpocapsae* survival during RD. We previously reported that *S. carpocapsae* IJ survival drastically declines below 64% RH, rendering them ineffective for infection (and therefore, biocontrol)^20^. Here, we assessed physiological responses, including rates of survival and water loss, in formulated and non-formulated IJs under low humidity conditions (64% and 54% RH), as well as the protection mechanisms provided by the formulations. We further investigated molecular strategies through transcriptomic analyses, comparing the results with those of previous slow-desiccation studies^16^ and *Heterorhabditis* data to understand the fundamental differences between slow and rapid desiccation. We then validated our novel findings through biochemical and ultrastructural approaches.

### Physiological basis of survival and protection mechanisms provided by formulations

We systematically characterized the physiological, biochemical, molecular and protective roles of two Pickering emulsions: TPE (oil-in-water [O/W]) and SPEG (W/O) during RD of IJs. *In-vitro* results demonstrated extended IJ survival through reduced water-loss rates, particularly below 64% RH, with survival prolonged to 144 h compared to 16 h in non-formulated controls. These findings, along with the observed variability in survival (post 72 h), highlight the protective efficacy of the formulations and indicate subtle yet significant temporal differences in their performance under RD stress. The observed survival differences correlated well with the gravimetric and FTIR data.

Specifically, FTIR analysis revealed distinct hydration dynamics for the TPE and SPEG formulations, with TPE exhibiting faster water loss compared to SPEG. These results were reinforced by a significant increase in efficacy with SPEG, as indicated in Ramakrishnan *et al*.^21^. Identified protective differences showed that TPE delays water loss, whereas SPEG retains the water phase. Furthermore, TPE also acts a system to compare as control for formulations without polymer. TPE is an O/W emulsion, with water as the major phase. Its protective capability arises primarily from a coating of oil and nanoparticles on the IJs, which reduces water permeability, supported by our epicuticle thickness results and consequently slows water loss, similar to the mechanism observed previously^33^. In contrast, SPEG is a W/O emulsion that protects IJs by maintaining hydration through a superabsorbent polymer layer, which may be further stabilized by an additional oil or nanoparticle layer, creating a layered protection system.

Biochemically, trehalose accumulation in formulated IJs increased consistently until 72 h of RD but remained lower than in non-formulated rapidly desiccated IJs. TPE and SPEG also exhibited contrasting accumulation patterns, further validating their distinct protective mechanisms and corresponding gene-expression trends. The variation in trehalose accumulation between the two emulsions confirms the protection mechanisms also supported by the cytoskeleton integrity observations. Both formulations preserved ECM integrity, thereby enabling survival of the EPN IJs. Hence, based on the above evidences, it can be concluded that, the survival benefits arise from a combination of several factors including the cuticle-nanoparticle interaction and hydration buffering.

### Glycocalyx enrichment and involvement of mucin-O-glycan pathway

Transcriptome analysis revealed that SPEG activates pathways similar to those activated in desiccated IJs but physiologically, resembles the response of IJs at 85% RH. Interestingly, mucin-O-glycan biosynthesis and glycosyltransferase were uniquely enriched in SPEG samples. Specifically, GalNac (N-acetylgalactosyltransferase; Tn antigen) encoded by *gly-7* showed significant enrichment. Moreover, several other genes related to this pathway exhibited FCs of at least 1.5 across all humidity conditions. Mucins are high-molecular-weight serine–threonine backbone-containing O-glycosylated proteins present in the cuticle and surface coat (glycocalyx) of many animal- and plant-parasitic nematodes^34, 35^. Several reports suggest their role in adhesion, hydrogel-forming abilities and protective properties^36^, conferring a negative charge to the cuticle due to their polyanionic nature^37^, while RNAi targeting of mucin genes leads to reduced adherence of endospores^35,38^. Although mucins have not previously been reported in Steinernematidae, wheat-germ agglutinin lectin-binding assays revealed carbohydrate moieties throughout the cuticle of *S. carpocapsae*, whereas they were restricted to the anterior tip in *H. bacteriophora* (Wray Carl Hansen III master’s thesis, Ment lab unpublished results;^59^), similar to observations from previous studies^40^ ^6445^. Proteins encoded by let-653 are structural constituents of the cuticle, and a mutation in *C. elegans* results in larval death ^46^. These proteins were significantly upregulated in SPEG (logFC: 3.1) and at 85% RH (logFC: 3.0) compared to RD at lower humidity (logFC: 2.5). Moreover, mucin-related genes were similarly upregulated under desiccation stress in *Ditylenchus destructor* ^25^, reinforcing their critical role in nematode desiccation tolerance. Based on TEM-observed glycocalyx enrichment, validating the transcriptomic observations, we propose that the highly negatively charged glycoproteins present in the glycocalyx attract the positively charged polymer layers in the formulations via electrostatic interactions, resulting in the retention of a water-rich layer on the IJ cuticle, as evidenced by changes in epicuticle thickness and sarcomere integrity.

### Rapidly desiccated vs. preconditioned IJs

Comparative analysis of enriched pathways and processes in preconditioned and rapidly desiccated IJs revealed that RD requires ∼50% more metabolic processes and pathways for adaptation, indicating a markedly high bioenergetic cost for the rapid stress response. A noteworthy finding was the similarity of processes activated between RD and later time points (36 h) of osmotic desiccation with PEG, where shared processes were marked by the upregulation of water-transport mechanisms. Further, the minimal overlap between evaporative desiccation and RD indicates that gradual physiological adaptations at higher humidity (97% RH) differ significantly from those elicited by rapid exposure to lower humidity (85% or 64% RH). Consequently, it explains the variations in phenotypic tolerance observed among EPNs^20^ and highlights the differences between various desiccation regimes. Interestingly, IRE-1-mediated UPR in the endoplasmic reticulum and Z-discs were shared by all types of desiccation conditions, indicating protein misfolding and aggregation are common. Furthermore, Z-discs are structural anchors that link the cuticle to the cytoskeleton and may act as critical signaling hubs^47,48^, potentially activating downstream protective effector proteins and pathways. Note that processes involving BM integrity and mechanosensory responses were enriched significantly for RD, highlighting a heightened physiological response to harsher conditions. Collectively, these findings suggest that RD represents a unique and highly demanding form of desiccation stress, characterized by high bioenergetic costs and extensive damage to structural and biomechanical integrity, which are crucial aspects in determining the organism’s survival capacity.

### Core pathways and strategies for survival under desiccation stress

A major desiccation-survival strategy involves the accumulation of hydrophilic proteins termed hydrophilins or IDPs^4,30^. During RD, we observed significant upregulation of LEA and DUR (Table S4), indicating that IDPs uniquely prevent or stabilize biomolecular entropy changes induced by desiccation, acting as molecular shields that prevent interactions. In contrast, tardigrades produce diverse groups of IDPs that stabilize cellular structures through gelation^49,50^. Despite significant gaps in our understanding of the physiological roles and cooperative mechanisms of these protective proteins, cytoskeletal reorganization and its potential role in preserving or stabilizing tissues, membranes and organelles during anhydrobiosis remain key areas for further investigation

### Trehalose accumulation does not guarantee survival

Trehalose acts as a chemical chaperone with high thermodynamic stability^51^ that protects membranes and proteins via “water-replacement theory” ^30,52,53^ and/or water entrapment and vitrification^24,49,54,55^. Intracellular accumulation of trehalose has been documented under various forms of stress. Alternatively, for quiescent organisms a diverse array of protectants have been reported among metazoans^56–58^. Bioprotection strategies vary among EPN species, for example, a preference for trehalose accumulation in *S. carpocapsae* compared to glycerol in *Steinernema feltiae* and *H. bacteriophora*^29,59^. Erkut and colleagues ^27,28,30^ suggested a minimum required threshold of trehalose accumulation during preconditioning in nematodes. Our findings indicate significant variations in trehalose accumulation over time. Correlating trehalose accumulation with our prior results of survival and water-loss rates ^20^, we concluded that the initial 6 h of RD are critical in delaying water loss, indicating that the timing of accumulation is crucial. During this window, EPNs could enhance permeability barriers, lipid packing and membrane stability, gaining time to activate protective mechanisms, including trehalose accumulation. Abusharkh *et al*.^60^ found that polar headgroups preferentially switch to phosphotidylethanolamine, facilitating trehalose insertion to replace hydrogen bonds. Our results show rapid trehalose accumulation at low humidity but also leads to low survival, indicating a lower protective effect of trehalose and probably suboptimal membrane stabilization. Conversely, at higher humidity, membrane fatty acids are surrounded by hydration shells providing optimal membrane adjustments for high survival, as depicted in Fig. 4E.

## Conclusion

The last decade of EPN research has focused on improving application and formulation systems^61^. We expanded upon this research by exploring EPN biology under RD conditions and examining the protective effects of Pickering emulsion formulations. Our findings demonstrate that RD is a unique form of desiccation stress, characterized by significant bioenergetic demands and substantial disruptions to mechanical stability in IJs. Using physiological, biochemical, molecular and ultrastructural analyses, we showed that RD adversely affects ECM integrity by disrupting BM assembly and cytoskeletal organization. Utilizing protected IJs as “gain-of-function” models, we confirmed that preserving ECM and sarcomere integrity by reducing water-loss rates is crucial for IJ survival and efficacy under RD conditions. This study thus provides fundamental insights into the biological mechanisms underpinning EPN tolerance and survival strategies during RD.

## Limitations of the study

The current study is limited mainly the genetic intractability of the nematode systems, that makes it difficult to extrapolate the regulatory and genetic role of the identified genes involved within the basement membrane and their impact on survival during rapid desiccation.

## METHODS

### EPNs

The original EPN populations were provided by e-nema GmbH (Schwentinental, Germany). *S. carpocapsae* ALL strain was cultivated in the last instar larva of *Galleria mellonella* at 23°C ^20^. The infected larvae were transferred to a modified White trap, and emerging IJs were collected and stored at 8°C until further use.

### Formulation treatments

SPEG was prepared from commercial hydrophobic silica (Aerosil R972, Evonik, Germany; fumed silica treated with dimethyldichlorosilane, estimated primary particle size of 16 nm) in paraffin oil and water (analytical grade; Sigma-Aldrich, St. Louis, MO, USA). Silica nanoparticles were dispersed in paraffin oil by sonication for 5 min (Sonics Vibra-Cell 750 W, 25% amplitude) with a silica content of 0.5 wt%. A solution of 0.5% potassium polyacrylic acid—0.5 g completely dissolved in 99.5 ml water by stirring for 1–2 h ^21^—was added at W/O ratio of 40:60 by volume and the mixture was sonicated for 10 min (at 25% amplitude) for emulsification. TPE was derived from amine-functionalized titania in water and mineral oil. The titania-NH_2_ nanoparticles were sonically dispersed in deionized water for 5 min at 1 wt%. Next, mineral oil was added at an O/W ratio of 4:6 by volume. The mixture was sonicated for 10 min at 25% amplitude to achieve emulsification^22^. *S. carpocapsae* IJs were vacuum-filtered, suspended in distilled water and adjusted to a concentration of 1000 nematodes/ml. The suspension was centrifuged at 4000*g* for 2 min and the pellet was vacuum-filtered. The IJ pellet was suspended in the required volume of formulation SPEG (W/O) or TPE (O/W) emulsion. This procedure was followed unless otherwise specified.

### Survival of formulated EPN *in vitro*

Survival of *S. carpocapsae* that had been rapidly desiccated at 64% or 54% RH was estimated and compared as described previously ^20^ with some modifications. To establish stable 64% RH, saturated calcium chloride salt and sodium nitrite were kept at opposite ends of the humidity chamber. This was done to stabilize the humidity at 64%RH, as the presence of formulations alone altered the humidity by +15-20%RH.Approximately 5000 vacuum-filtered IJs were suspended in 100 µl treatment solution (water, TPE or SPEG), spotted on Teflon discs and dried in a laminar airflow chamber for 20–45 min to remove excess water. Samples were transferred to a desiccator at 64% or 54% RH maintained at 23°C with continuous humidity-monitoring by data loggers (LOG32TH and SSN23E). Samples were collected at 0 (immediately after drying), 16, 24, 48, 72, 96, 120 and 144 h of RD. Samples were excised from the disc, suspended in distilled water and incubated overnight to assess nematode survival rate by randomly counting the number of live or dead IJs out of 200 IJs under a binocular (Olympus SZH10, 30×lJmagnification). IJs were scored as live based on active movement in response to probing. Data were used to calculate survival rate as percentage of total number of nematodes. The experiment was repeated three times with five replicates per treatment.

### Evaluating EPN water content by gravimetric method

Water content of EPNs during RD was estimated using the gravimetric method described by Ramakrishnan *et al.* (*34*) with slight modifications. Water or emulsions (120–150 µl) were drop-casted on slides with Teflon sheets, and IJs (25,000–50,000) were vacuum-filtered and gently transferred to the slides. This prevented any loading errors for IJs and emulsions. The samples were dried under laminar airflow for 20 min, and set as the 0 h samples; these were weighed and transferred to desiccators maintained at the specified RHs (64 ± 5% or 54 ± 4%). Control treatments consisted of 120–150 µl of water or emulsions that were similarly processed. All samples and controls were weighed at 24 h. Gravimetric measurements post-24 h were not reliable due to negative values obtained for SPEG samples when the negative control (SPEG emulsion without IJs) was subtracted. Dry weight was estimated by oven-drying at 90 or 105°C for 12 h or 3 h, respectively. Water weight was calculated by subtracting the dry weight of the nematode samples from the total estimated weight at different time points. The weight of the control samples was also deducted to remove the effect of residual water or emulsions on water loss. Hence, the water content of nematodes estimated as weight (mg) was expressed as a percentage (%), i.e., normalized body water content, using the formula:

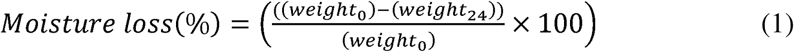

where, weight_0_ is the initial sample weight at 0 h and weight_24_ is the sample weight at 24 h.

### Statistical analysis

Data from repeated experiments were combined to analyze the differences in formulated vs. control IJ survival. Survival data were arcsine-transformed and subjected to ANOVA, and significant differences among pairs were further elucidated through Tukey-HSD or Student’s t-test comparisons. Water-loss measurements were subjected to two-way ANOVA comparisons. Results for trehalose accumulation in rapidly desiccated IJs were subjected to ANOVA and test slices were carried out for post hoc comparison by humidity and time point. Trehalose accumulation at 0 h was analyzed by one-way ANOVA followed by Tukey multiple comparisons. Trehalose accumulation for formulated IJs was subjected to ANOVA and significant differences among each pair were further elucidated through Tukey-HSD or Student’s t-test comparisons. Data related to cuticle and epicuticle thickness and annulation spacing were tested by one-way ANOVA with means and compared using Tukey multiple comparison post-hoc. All comparisons were carried out with JMP statistical software.

## Data Availability

The authors confirm that the data supporting the findings of this study are available within the article and its supplementary material.

## Author Contribution

All authors contributed to the study’s conception and design. JR wrote the first draft of the manuscript and all authors commented on versions of the manuscript. Conceptualization: JR, DM, GM, DSI; Methodology: JR, DM, IG, GM, SSV, AN, MS, EB, VR, KAM; Data curation: JR, SSV, AF, AH, MS, AN, EB; Formal analysis and investigation: JR, SSV, AF, KAM; Writing—original draft preparation: JR, DM; Writing—review and editing: JR, DM, IG, VR, SSV, KAM, GM, DSI; Funding acquisition: DM, GM, DSI; Resources: DM, AF, AN, GM. Supervision: DM, IG, GM. All authors have read and agreed to the published version of the manuscript.

## Acknowledgements

We would like to thank Reut Amar Feldbaum and Michael Brichka for providing the emulsions, and Dr. Einat Zelinger and Dr. Tally Kossovsky for their assistance with cryo-TEM sample preparation and imaging. Our heartfelt gratitude goes to Dr. Hillary Voet for statistical consultations. This study was partially funded by the Chief Scientist of the Israeli Ministry of Agriculture (project number 20-06-0085) and US-Israel Binational Agricultural Research and Development Fund (BARD IS-5183-19).

## Notes

### Competing Interest Statement

The authors have declared no competing interest.

### Summary of Updates

methodologies was updated based on suggestions

